# Spike: A GPU Optimised Spiking Neural Network Simulator

**DOI:** 10.1101/461160

**Authors:** Nasir Ahmad, James B. Isbister, Toby St. Clere Smithe, Simon M. Stringer

**Author notes:** Correspondence: Nasir Ahmad.

## Abstract

Spiking Neural Network (SNN) simulations require internal variables – such as the membrane voltages of individual neurons and their synaptic inputs – to be updated on a sub-millisecond resolution. As a result, a single second of simulation time requires many thousands of update calculations per neuron. Furthermore, increases in the scale of SNN models have, accordingly, led to manyfold increases in the runtime of SNN simulations. Existing solutions to this problem of scale include high performance CPU based simulators capable of multithreaded execution (“CPU parallelism”). More recent GPU based simulators have emerged, which aim to utilise GPU parallelism for SNN execution. We have identified several key speedups, which give GPU based simulators up to an order of magnitude performance increase over CPU based simulators on several benchmarks. We present the Spike simulator with three key optimisations: timestep grouping, active synapse grouping, and delay insensitivity. Combined, these optimisations massively increase the speed of executing a SNN simulation and produce a simulator which is, on a single machine, faster than currently available simulators.

## 1 INTRODUCTION

Point neuron based Spiking Neural Networks (SNNs) have been the focus of scientific investigation for several decades. These kinds of models may be used to simulate the ‘spiking’ dynamics of real neurons in the brain, which communicate with each other by emitting electrical pulses called action potentials or ‘spikes’. In this pursuit, a number of SNN simulators have emerged (Vitay et al., 2015; Zenke and Gerstner, 2014; Stimberg et al., 2014; Yavuz et al., 2016; Linssen et al., 2018), which provide combinations of high speed execution, hardware support, and simplicity when defining models.

The simulation speed of SNNs is particularly difficult to optimise. This is due to both the neuron temporal dynamics and their interruption by incoming synaptic events. Spiking neurons have at minimum a single internal variable referencing their membrane voltage. Commonly, the dynamics of this membrane voltage are solved numerically at a sub-millisecond timestep. Though there are alternative, sometimes analytic, methods to solve these dynamics, in a network of interacting neurons any numerical integration method requires interruption upon the arrival of spikes to a neuron. This interruption is necessary due to both the discontinuous nature of neuron membrane voltages during an action potential, and the discontinuous synaptic input changes upon the arrival of a pre-synaptic spike. Upon each of these events, the membrane dynamics are altered in a non-integratable fashion and therefore even analytic or higher-order numerical integration schemes must halt at the discontinuity and be restarted. For this reason, the forward-euler first order numerical integration method is commonly used for SNN simulations, which means that small timesteps are required.

Networks require many thousands of membrane voltage updates for a single second of simulation time. This is a large number of updates which we must consider before even addressing the number of calculations required to deal with synaptic inputs or synaptic learning. Moreover, in many networks, the number of synapses far exceeds the number of neurons, and so these synaptic computations entail even greater computational complexity.

Just as GPU hardware developments have brought major performance boosts to fields such as image processing, deep learning, and data processing at scale, they have also been used for SNN simulation. However, a number of GPU SNN bottlenecks remain unresolved. This paper proposes effective solutions to a set of GPU SNN bottlenecks and presents these in a new GPU based SNN simulator, Spike.

Finally, the significant benefits of GPU based simulators over existing CPU based simulators are yet to be compared in like-for-like benchmarks. We therefore extend a set of benchmarks to GPU-based simulators and make available a repository in which all code used to produce these benchmarks is open for continued access and evaluation by the community.

## 2 METHODS

### 2.1 Background

Applications which leverage GPU devices make use of a paradigm in which procedures to be executed on data are defined as a “kernel”. Kernels, when run, process each element of the data in parallel over many processing threads on the GPU device. In SNN simulations, these kernels are defined to carry out actions such as updating the neuron membrane voltage and converting neuron spikes into the appropriate synaptic inputs.

This results in a simulation style in which the neuron update kernels are traditionally run for every timestep of the simulation, of which there may be many tens of thousands for a simulation of a few seconds. As the dynamics of the network must be synchronised in time, the host must wait until all kernels are complete before progressing to the next timestep. This process of repeatedly running the same kernel upon the same data is different to other data processing algorithms run on GPUs in which single kernels are often only run once upon a single set of data. The action of repeatedly executing kernels on each timestep exacerbates an overhead commonly referred to as a “kernel launch overhead”. Furthermore, each kernel launch computes a minimal amount of work, only processing spikes from a single timestep and thereby reduces the efficiency of the kernels further. In order to reduce the impact of the kernel launch overhead and to maximise the amount of work per kernel launch, we propose a novel technique we refer to as Timestep Grouping (TG).

Significant computational time can also be lost when a kernel is launched and not all threads are required. The occupancy of a kernel is a term used to refer to how many of the threads are carrying out computations. A major issue in SNN simulations is that on a given timestep, most (if not all) synapses have no spikes arriving. This results in the kernels dealing with synapses having a very low occupancy. We describe and test several methods for overcoming this issue of occupancy and ultimately identify a technique hereafter referred to as Active Synapse Grouping (ASG). This technique is leveraged by many existing SNN simulators but has yet to be formally published in comparison to other approaches.

We wished to highlight ASG in order to also compare it to recent suggestions for alternative synaptic processing such as the use of Dynamic Parallelism (Linssen et al., 2018) and to show the combined efficacy of TG and ASG in a GPU based SNN simulator. Though ASG is also implemented by the GPU based SNN simulator GeNN (Yavuz et al., 2016), the introduction of TG and the combined use of ASG and TG on a GPU platform is a novel feature of Spike.

### 2.2 Timestep Grouping

On a GPU device, the execution of a kernel causes a small delay, previously referred to as the kernel launch overhead. This delay is compounded over the course of a SNN simulation by the sheer number of kernel launches, eventually leading to a significant computational cost.

In order to mitigate this issue, we leverage a feature of many SNN models; delayed propagation of spikes from neuron to synapses. Many SNN models define individual synapses as having some delay in the arrival time of a spike from the pre-synaptic neuron. This is often referred to as an axonal delay. We leverage this feature to implement timestep grouping.

Timestep grouping requires the determination of the smallest axonal delay in the network. Thereafter, the network’s smallest axonal delay is used to update the network in grouped computations. Kernels are launched, updating the network dynamics at the numerical timestep by way of loops as shown in Figure 1. Figure 1 A shows the regular timestep based update in which each the network state is updated via a kernel launch upon each timestep. Timestep grouping allows a single kernel to update the network multiple times (up to the timestep grouping value) as described in Figure 1 B. This timestep grouping is determined by calculating the size of the smallest axonal delay in the network as measured in the number of timesteps.

**Figure 1.**
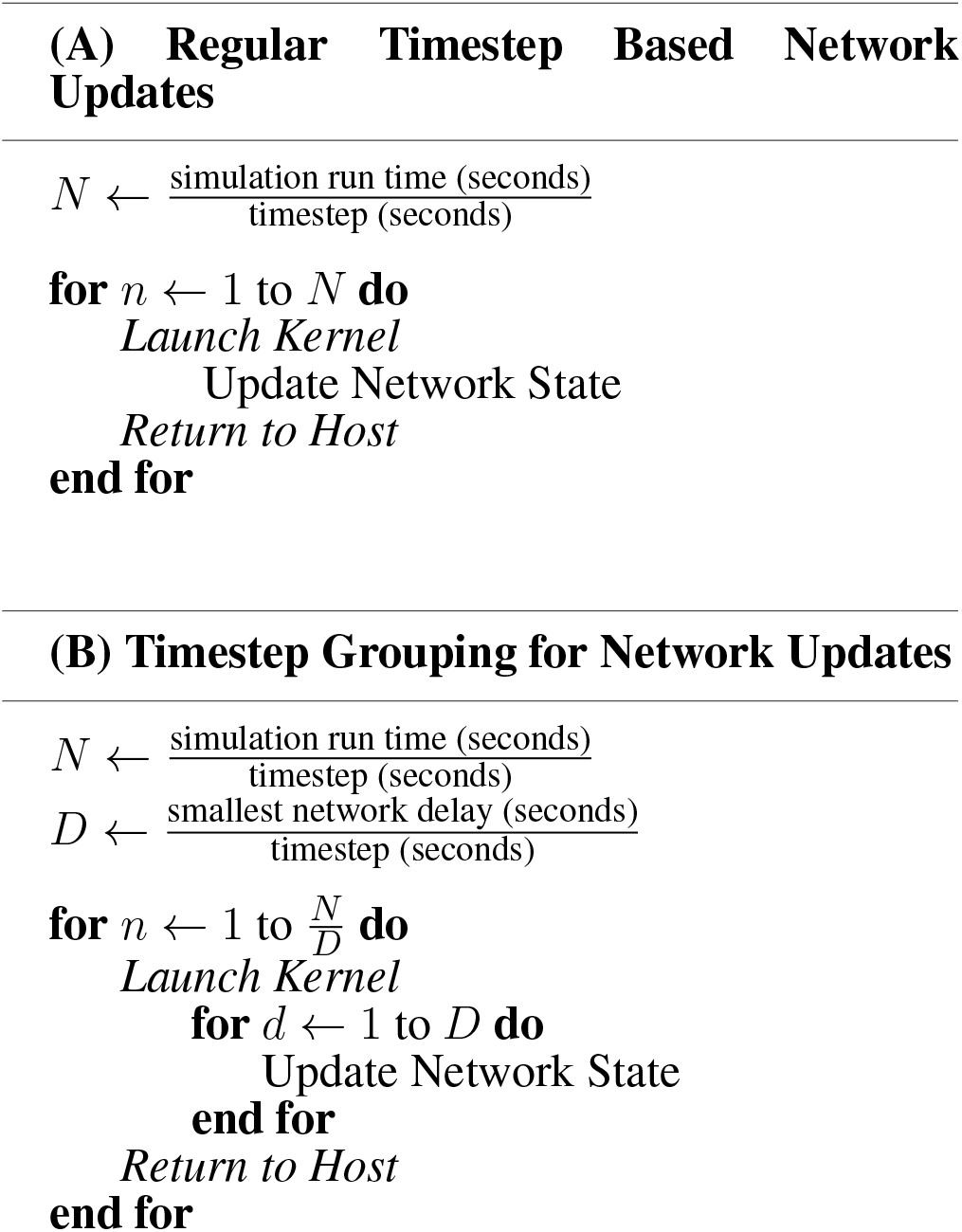
Network Update Optimisation. **(A)** Looping structured used for regular timestep based network updates. **(B)** Looping structure used for timestep grouping.

Any optimisations must ensure that the modelled SNN dynamics are unaffected. For timestep grouping we leverage the fact that for any time period of the simulation which is smaller than the minimum axonal delay, only spikes which were emitted before that time period can affect neuron dynamics during that time period (given the nature of delays). Therefore, if timestep grouping is kept smaller than (or equal to) the minimum axonal delay it avoids affecting network dynamics by continuing to integrate activity which was recorded before the beginning of the timestep grouping. Furthermore, any spiking activity within this period only affects other neurons after the minimum axonal delay and can therefore be collected at the end of the timestep grouping for use in the next timestep grouping.

In GPU based computation, a given kernel is run many times in parallel and there is no guarantee on the order in which these parallel threads are run. For example, if ten neurons are updated with one kernel launch, we cannot be sure as to which neuron will be updated first. However, since the order of the update within a timestep grouping period does not affect dynamics we can ignore this lack of synchronised computation. Instead computation is synchronised after each timestep grouping and all network activity recorded. This synchronisation of computations is what is referred to in Figure 1 as “Return to Host” as all host-side computations are executed in series. With an appropriate timestep grouping (i.e. smaller than or equal to the minimum delay) the neuron dynamics are thereby kept consistent while the number of kernel launches are reduced by a factor equal to this timestep grouping.

A further benefit of the timestep grouping is the greater amount of work per kernel call. Since kernels computing network updates require synchronization after each update, the time required for a single update to the network is bound by the slowest thread. Taking neurons as an example, the slowest threads would be those on which the neuron fires an action potential and requires its membrane voltage to be reset and an action potential to be propagated. Spiking events for a single neuron are sparse on the timescale of a timestep meaning that for timestep based kernel executions, the majority of neurons must await the excess computations for the few neurons which have spiked. In contrast, timestep grouping results in the grouping of computations for a number of timesteps. Thus, a greater number of action potentials can simultaneously be processed in a single kernel execution and more processing threads leveraged.

The reduced kernel launch overhead and greater work per kernel execution ultimately reduces simulation time, as explored in the Results section below.

### 2.3 Active Synapse Grouping

The number of synapses in SNN models often exceed the number of neurons by orders of magnitude. The synapses have a few parameters specific to their function, including the synaptic weight, and most often contribute to either instantaneous current injections or to the excitatory/inhibitory conductances of the post-synaptic neurons. However, the synapses only require processing when a spike from a pre-synaptic neuron reaches them (after an axonal delay).

Checking all of the synaptic connections on every timestep is the most algorithmically simple method for determining when spikes due to recent firing of pre-synaptic neurons should be propagated (shown in Figure 2 (A)). This results, however, in a large number of threads being launched which check synapses that require no action due to the inactivity of the pre-synaptic neuron. These threads contribute to a significant inefficiency in computation.

**Figure 2.**
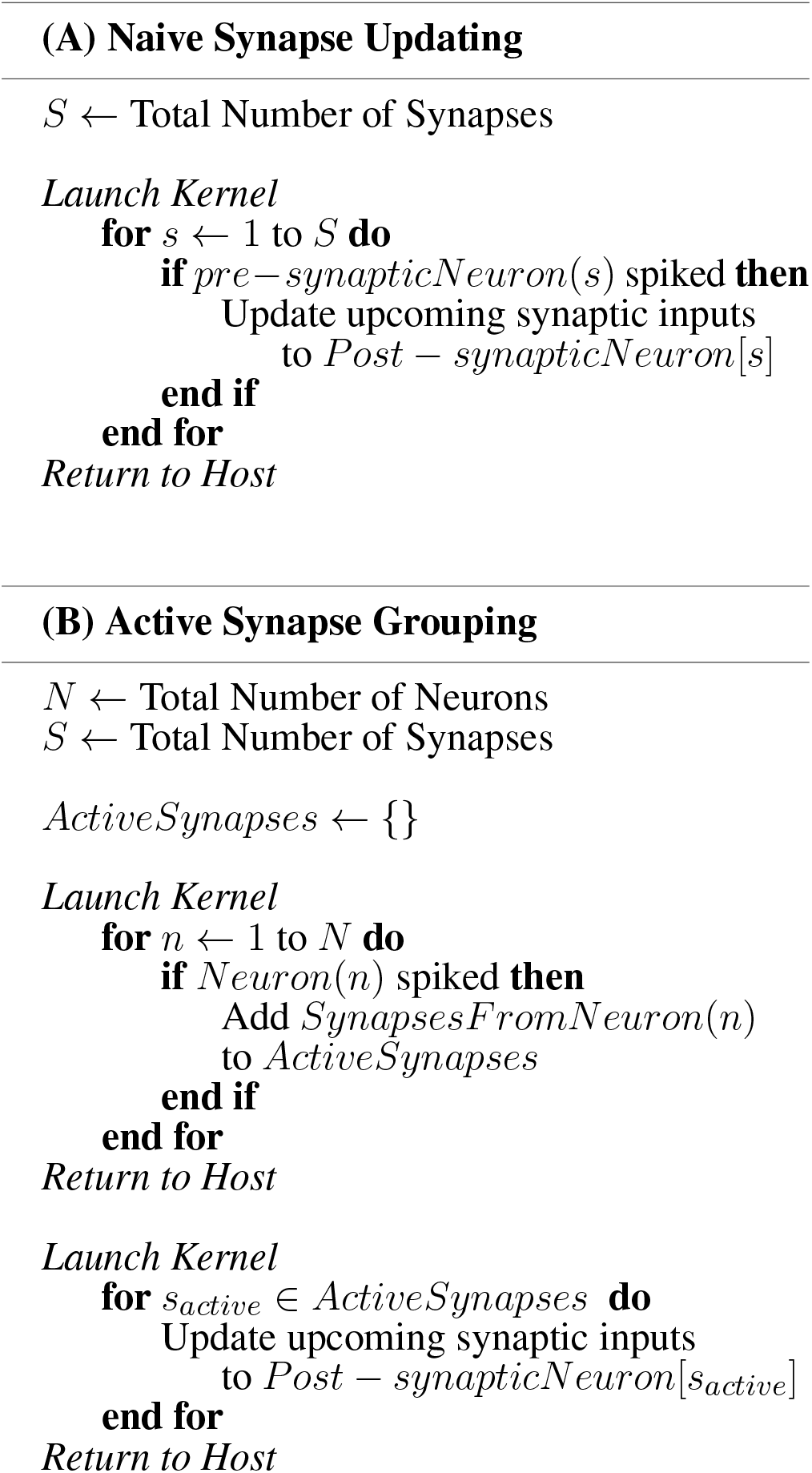
Synaptic Update Optimisations. **(A)** The computationally most simple method for detecting synaptic activations. **(B)** A description of the two stage process required for Active Synapse Grouping (ASG).

Active Synapse Grouping (ASG) is an algorithmic alternative to this approach. In this approach, a neuron kernel is launched which loops through all of the neurons to check if they have emitted a spike. If a neuron has spiked, then its efferent synaptic connections are flagged as active. After the neuron kernel is complete, a second kernel is launched in order to update only the flagged synaptic connections. Thus, only active synaptic connections are considered for computation. This process is outlined in Figure 2 (B). Axonal transmission delays would correspond in this approach to an adjustment during the updating of active synaptic connections. We implement a buffer based approach which is outlined in Section 2.4.1. Ultimately, synapses can be dealt with efficiently and computation is kept to a minimum.

The effect of ASG on simulation time is explored in the results section below, where it is compared to a basic synapse update algorithm and a recently proposed GPU specific optimisation, referred to here as Dynamic Synapse Parallelism (DSP) (Kasap and van Opstal, 2018).

### 2.4 The Spike Simulator

The Spike simulator has been under development since 2015 with the express aim of producing a high speed GPU based SNN simulator. It is written in C++ and CUDA with a flexible hierarchical class structure for Neuron, Synapse, and Plasticity Models. The past two sections describe optimisations (TG and ASG), which have been implemented in the Spike Simulator. Another key goal of the Spike simulator development was to be computationally efficient when dealing with delayed synaptic transmission between neurons. In particular, we aimed to ensure that no overhead was brought to a simulation even if synaptic connections have heterogeneous distributions of axonal delays. This required some thought to ensure that synaptic processing was not interrupted or made more complex.

#### 2.4.1 Delay Insensitive

A simulator whose performance is unaffected by axonal delays is particularly desirable given the range of research avenues which leverage delays within spiking neural networks. SNN models investigating a range of subjects including sound localization, reservoir computing, auditory processing and visual feature binding (Goodman and Brette, 2010; Paugam-Moisy et al., 2008; Erfanian Saeedi et al., 2016; Eguchi et al., 2018) are but a few of the many studies which implement multiple axonal delay values within a single study for modelling purposes. Spike is a simulator designed to optimise this process.

In order to meet the aim of a delay insensitive simulator, every neuron in a simulation is assigned a set of input buffers (one for each synapse type, e.g. excitatory and inhibitory). The neurons iterate over these buffers to collect incoming synaptic updates. Upon each timestep, a single neurons shifts forward on its buffers by one location and reads the synaptic inputs for its current timestep. These buffers are circular, meaning that upon reaching the end of a buffer calculations return to the beginning again, and therefore the buffer can be treated as a ring.

If we define the length of this buffer as equal to the maximum possible synaptic delay, synaptic inputs can be placed in this buffer at a distance (corresponding to the delay of that synapse) from where the neuron is currently querying for inputs. This distance would correspond to the synaptic delay in timesteps and would therefore ensure that the neuron will only reach this input after the corresponding delay. The circular nature of the buffer ensures that memory is conserved and since no incoming spike affects the neuron at any time delay longer than the maximum delay, it is sized efficiently.

Since no excess calculations are necessary, the simulation speed is unaffected by the inclusion of any delay structure in the synaptic transmission. The only consideration to be made is that of memory (these buffers introduce a memory overhead per neuron) though the Spike simulator is focused upon speed rather than memory efficiency.

### 2.5 Benchmarks

#### 2.5.1 Network Models

Simulators are compared in this paper using two previously published benchmarks hereafter referred to as the Vogels-Abbott and Brunel benchmarks (Vitay et al., 2015; Zenke and Gerstner, 2014). The Vogels-Abbott benchmark is based upon a reduced scale version of the network presented in the Vogels and Abbott (2005) publication. This network consists of 3200 excitatory and 800 inhibitory Leaky Integrate and Fire (LIF) neurons with a 2% random synaptic connectivity. Synaptic connections follow a conductance based current input model and the network, driven by background stimulation, maintains a firing rate close to 17Hz. The specifics of the network dynamics and parameters are detailed in Appendix A.

The second benchmark network is one designed to test a larger network with a higher degree of connectivity. The Brunel benchmark is based upon a network adapted by Zenke and Gerstner (2014), published in its original form by Brunel (2000). It consists of 10,000 Poisson firing input neurons exciting 8,000 excitatory and 2,000 inhibitory LIF neurons. These LIF neurons have a 10% random connectivity with voltage injecting synapses. Appendix B details the dynamics of this network.

Furthermore, the Brunel benchmark network has two modes; with and without plasticity. The plastic mode consists of a weight-dependent STDP rule operating on the excitatory to excitatory LIF Neuron connections. This STDP rule is expected to lead to a normal weight distribution with a mean of 0.1mV. The details of the STDP rule are explained in Appendix B.1.

The benchmark network parameters are equivalent to those implemented by Zenke and Gerstner (2014) and Vitay et al. (2015). The code used to produce the comparisons against other simulators (below) have been collected in the following repository:

https://tinyurl.com/y7gltmsw

#### 2.5.2 Simulator Versions

The simulators compared in this study include four CPU based simulators – ANNarchy, Auryn, Brian2, and NEST (Vitay et al., 2015; Zenke and Gerstner, 2014; Stimberg et al., 2014; Linssen et al., 2018) – and a single GPU based simulator – GeNN (Yavuz et al., 2016).

Simulators were collected for comparison and added to the repository (detailed below) prior to September 2018 from each of their git repository master branches. The specific versions of the simulators compared can be viewed at the repository listed above, or can be determined using the following git commit IDs for each simulator:

ANNarchy @ 3f0e1d2
Auryn @ 6928b97
Brian2 @ 3ded00d
GeNN @ a4387e5
NEST @ d175510
Spike @ 9fd6235

#### 2.5.3 Test System

All single thread benchmarks described in this paper were produced in a system with the following specifications;

- CPU: Intel i7-4770K
- GPU: NVIDIA GTX 1070 founders edition

The system used to test the multithreaded performance has specification;

- CPU: 2x Intel Xeon E5-2600 v4 (16 cores, 32 Hyperthreads combined)

## 3 RESULTS

### 3.1 Vogels-Abbott Benchmark

This benchmark, as described in Section 2.5.1 and Appendix A, consists of 4000 Leaky-Integrate and Fire (LIF) neurons (3200 excitatory and 800 inhibitory) with a 2% random connectivity on conductance based synapses. This random connectivity was produced once and thereafter loaded for all results and simulators shown below. The neurons in the network are excited by a background 200pA input current (20mV per second) and produce chaotic dynamics under the network connectivity. This benchmark is used to describe the relative impacts of the various optimisations described above and to place the speed of the Spike simulator in the context of other available simulators.

#### 3.1.1 The Spike Simulator and Optimisations

The efficacies of the optimisations described in the Methods sections 2.2 and 2.3 are hereafter tested in the context of the Vogels-Abbott benchmark.

Figure 3 compares the simulation time required for the Vogels-Abbott benchmark in the Spike simulator across a range of conditions. It compares no optimisation (NONE) in which the synaptic updates are carried out individually in a naive fashion to a range of optimised conditions. In the NONE case, on each timestep every synapse tests whether its pre-synaptic neuron has fired and thereafter carries out any synaptic updates. This condition is compared to two other synapse relevant optimisations; Active Synapse Grouping (ASG), as described in the methods section, and Dynamic Synapse Parallelism (DSP). DSP refers to an approach which makes use of a fairly recent addition to the capability of NVIDIA GPUs in which individual threads on the GPU device can, themselves, launch more kernels (recruiting parallel threads). This is called Dynamic Parallelism. Dynamic Parallelism was identified by Kasap and van Opstal (2018) as a potential speedup when compared to a limited set of other synaptic update methods.

**Figure 3.**
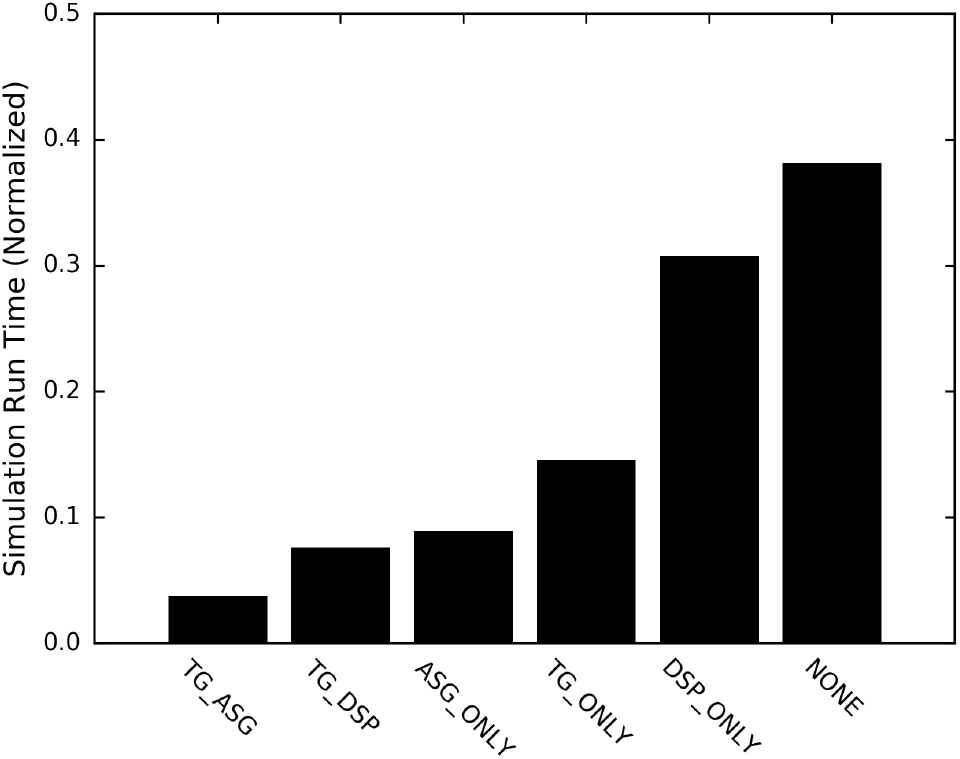
The effects of various optimisations upon GPU SNN performance. Comparing the speed of simulations in the Spike simulator under a range of optimisation conditions. NONE; No optimisations, ASG; Active Synapse Grouping, DSP; Dynamic Synapse Parallelism, and TG; Timestep Grouping. These optimisations are also shown in combinations. The optimisations are presented fastest first (shorter is faster).

Other than synaptic optimisations, we also compare the simulator speed with and without the inclusion of Timestep Grouping (TG). TG is further combined with either ASG or DSP in order to show the speedup of combining these optimisations.

Figure 3 shows a clear speedup under all optimisation conditions compared to the non-optimised case (NONE). The optimisation comparison also shows that ASG provides a significantly greater speedup than DSP. This is attributed to the fact that even though DSP ensures that non-active synapses are ignored, it nonetheless requires device-side kernel launching proportional to the number of spikes emitted during the simulation. Thus when compared to ASG (which only requires a single kernel launch per timestep and deals exclusively with active synapses) DSP is markedly slower. This furthermore indicates that the comparison carried out by Kasap and van Opstal (2018) should be revisited with ASG as another case for comparison.

TG shows a significant speed increase both in the presence of synaptic optimisations and without. Since the network computation is identical under the NONE and TG cases, the TG benefit is attributed to the reduction of the kernel launch overhead and the increased work per kernel launch (see Methods 2.2). In the Vogels-Abbott benchmark, the minimum axonal network delay is eight times larger than the network timestep and therefore the number of kernel launches is reduced eightfold and the number of spikes processed per kernel launch eight times higher on average.

#### 3.1.2 Comparing Simulators (Single Threaded)

Having established the best combination of optimisations in the previous section, we now compare the Spike simulator (with the TG and ASG optimisations) to a set of competing GPU and CPU simulators. Note that the CPU based simulators can simulate networks with multiple CPU threads, however we first consider all simulators with a single CPU thread.

Figure 4 compares the normalized simulation time required for the Vogels-Abbott benchmark by a set of SNN simulators. Hereafter, normalized simulation time indicates the average time taken per second of a simulation, when averaged over simulations of length 100 seconds. The simulators are ordered by speed. Note that shorter bars here indicate a faster speed (shorter is better) and that the y-axis is logarithmically scaled.

**Figure 4.**
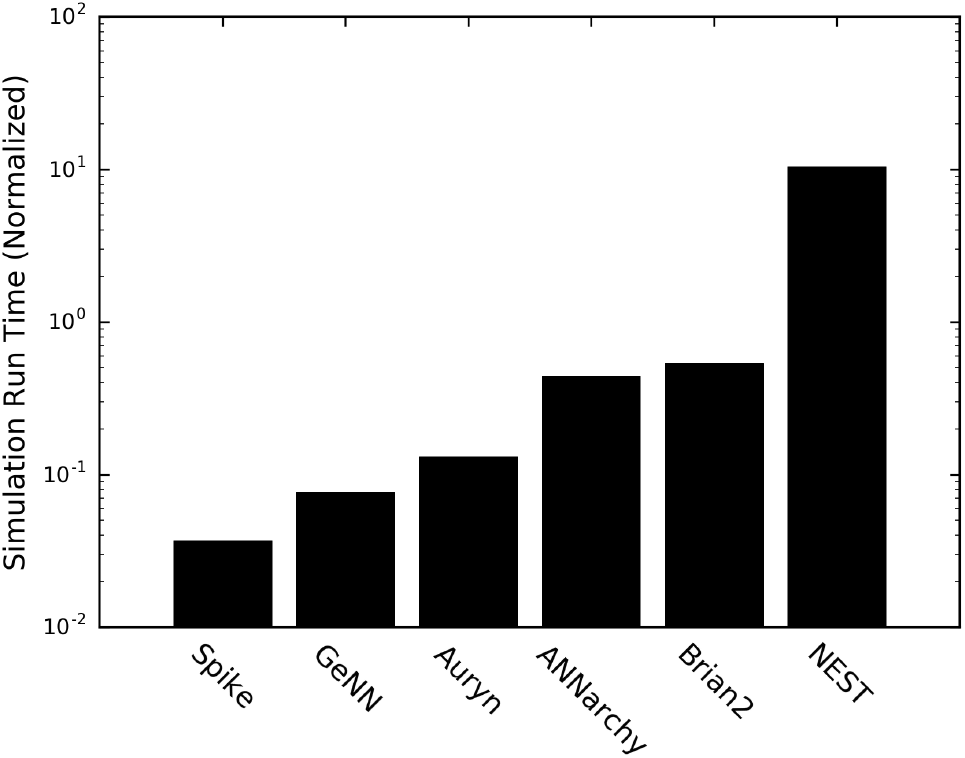
The Vogels-Abbott benchmark comparison across simulators. All single threaded or single GPU. Comparing the speed of a range of available simulators on the Vogels-Abbott benchmark. Shorter bars are faster. Note the log scale on the y-axis. These are ordered in speed, fastest first.

Figure 4 shows the benefits of GPU based simulators generally but also the particular simulation efficiency of the Spike simulator. Both GeNN and Spike (the only GPU based simulators in this list) show a significant speedup compared to competing simulators. These simulators are closely followed by Auryn which has been reported as the fastest simulator since its release (Zenke and Gerstner, 2014; Vitay et al., 2015). This benchmark shows a like-for-like comparison of these simulators and their relative speeds. The NEST is an exception on this list as it is the only simulator shown here which does not update the neuron dynamics with a forward-euler solver and instead makes use of a higher-order Runge Kutta solver. Previous studies (Zenke and Gerstner, 2014) have shown that when the NEST is modified to simulate LIF neuron dynamics with a forward euler solver, it gains significant ground in simulation speed. However, not only are the patches used for such a modification to the NEST out of date, the same paper also showed that nevertheless, the Auryn simulator remains the fastest CPU based SNN simulator. Thus, we use the performance of Auryn as a bound upon existing CPU based simulator speed.

#### 3.1.3 Multithreaded CPU Performance

Comparison of a single GPU versus a single threaded CPU based simulation ignores the performance of multithreaded CPU based simulators. Given the multithreaded benchmark results presented by Zenke and Gerstner (2014) and Vitay et al. (2015), Auryn remains the leading CPU based simulator both in the single and multithreaded use case. We therefore use Auryn as a comparison simulator.

Figure 5 compares the speed of Auryn in multithreaded execution mode to the Spike simulator on a single GPU. To date, no GPU based simulators, including Spike, appear to support multi-GPU computation and therefore the speed referenced here for Spike is identical to that shown in Figure 4.

As can be observed, only under multithreading with eight threads does the Auryn simulation approach the speed of Spike on a single GPU.

**Figure 5.**
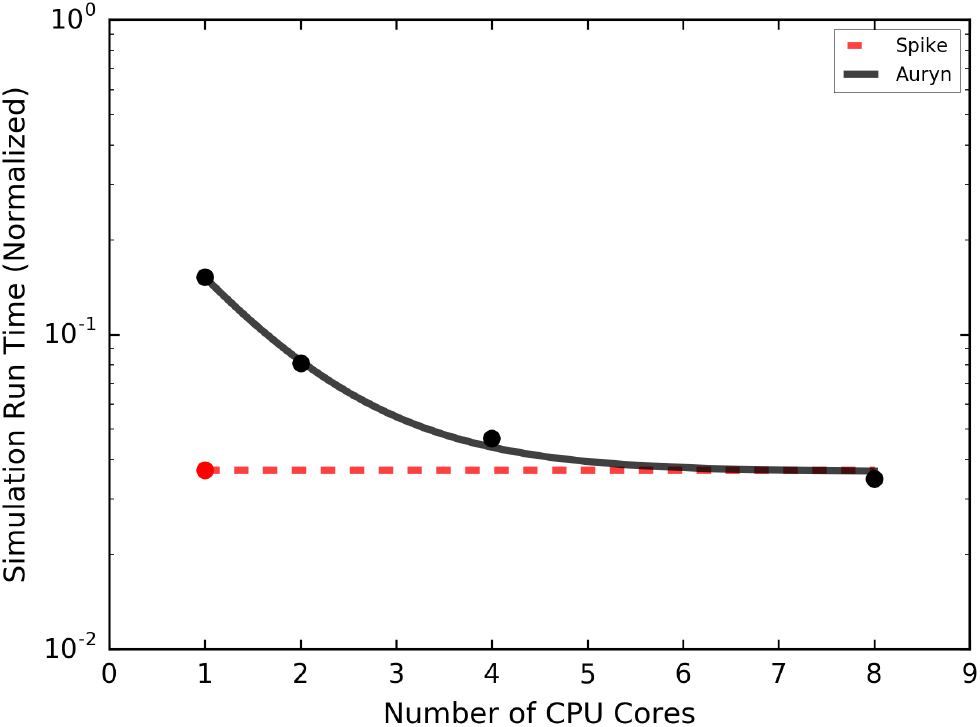
Comparing multithreaded CPU performance to Spike on a single GPU. This plot shows Spike in a single GPU mode only (therefore the dotted line is not a prediction of it’s performance over more CPU cores but instead just a baseline). Auryn was benchmarked with 1, 2, 4 and 8 CPU cores. An exponential decay curve was then fit to these points to produce the plot above.

#### 3.1.4 Simulating with multiple axonal delay values

As described in the methods section, one focus of the Spike simulator was to allow any delay structure in the synaptic connectivity. This feature of the Spike simulator is not shared by many other simulators. The Auryn, ANNarchy, and GeNN simulators do not, in their current states, allow heterogeneous distributions of axonal delays on synaptic connections. Instead, a user can create multiple synaptic populations, each with a different but homogeneous delay.

Figure 6 shows a comparison of the Spike simulator to the closest competing GPU simulator, GeNN (Yavuz et al., 2016) (see Figure 4). The simulators are compared in cases where the excitatory-to-excitatory connectivity has either a homogeneous delay on all synapses or a set of synaptic populations where each population has a unique delay value. In the Spike simulator, this is achieved without the need for multiple synaptic populations. Instead, a single synaptic population can have any delay structure. GeNN (alongside Auryn and ANNarchy) requires the creation of multiple synaptic groups.

**Figure 6.**
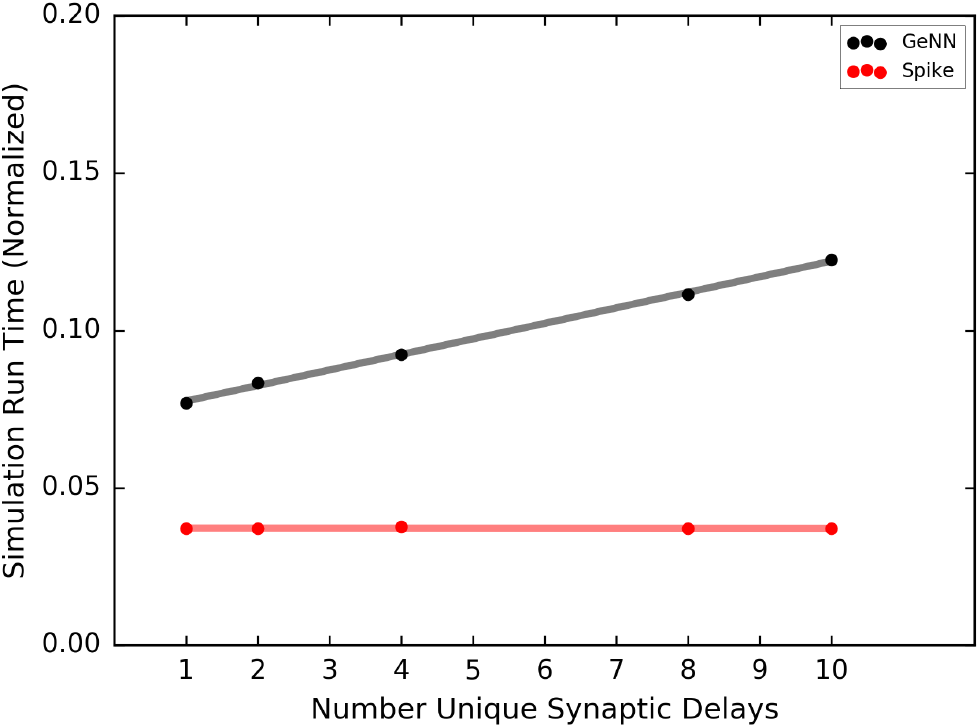
Comparing the effect of adding a range of delayed synapses upon simulation speed in Spike versus GeNN. The simulation speed is shown for a variant of the Vogels-Abbott benchmark in which the synaptic delays in the excitatory to excitatory connections is varied between either homogeneous or multiple sets of synapses each with a unique delay for their group.

As can be seen in Figure 6, Spike produces no change in simulation speed with changing delay structure. By comparison, the increasing number of synaptic populations required by GeNN result in a linear slow down of the simulation. The Vogels-Abbott benchmark is particularly suited to this comparison as the introduction of axonal delays to the synaptic connectivity does not affect the firing rate in a significant manner.

This comparison highlights how well the Spike simulator reaches its goal of a delay accommodating simulator. Synaptic connections which a neuron makes to itself (autapses), multiple connections between a pairs of neurons (multapses), and synaptic connections with different delays are treated no differently in the simulation framework from any other synaptic connection and do not contribute to any change in the simulation speed (assuming the network average firing rate is unaffected).

### 3.2 Brunel Plasticity Benchmark

A second SNN benchmark used to compare the simulators above is the Brunel benchmark consisting of 10,000 Poisson firing input neurons, and 10,000 LIF neurons (8,000 excitatory and 2,000 inhibitory) as detailed in Section 2.5.1 and Appendix B. This network is more dense than the Vogels-Abbott benchmark with a 10% connectivity between the LIF neurons and a 10% connectivity from the Poisson neurons to the LIF population. This benchmark therefore represents a significantly larger and more densely connected network than the Vogels-Abbott network. Furthermore, a weight dependent STDP rule is used to update the excitatory to excitatory synapses as described in Appendix B.1. This allows a second comparison between the SNN simulators to test their efficiencies when computing weight updates through synaptic plasticity.

Figure 7 shows (as above for the Vogels-Abbott Benchmark) a comparison of the normalized simulation time for the range of simulators with the Brunel benchmark. The black and red bars show normalized simulation time for the same network with and without plasticity running respectively. As with the Vogels-Abbott benchmark, these results are based upon the simulators loading a network with identical connections between the LIF neurons.

**Figure 7.**
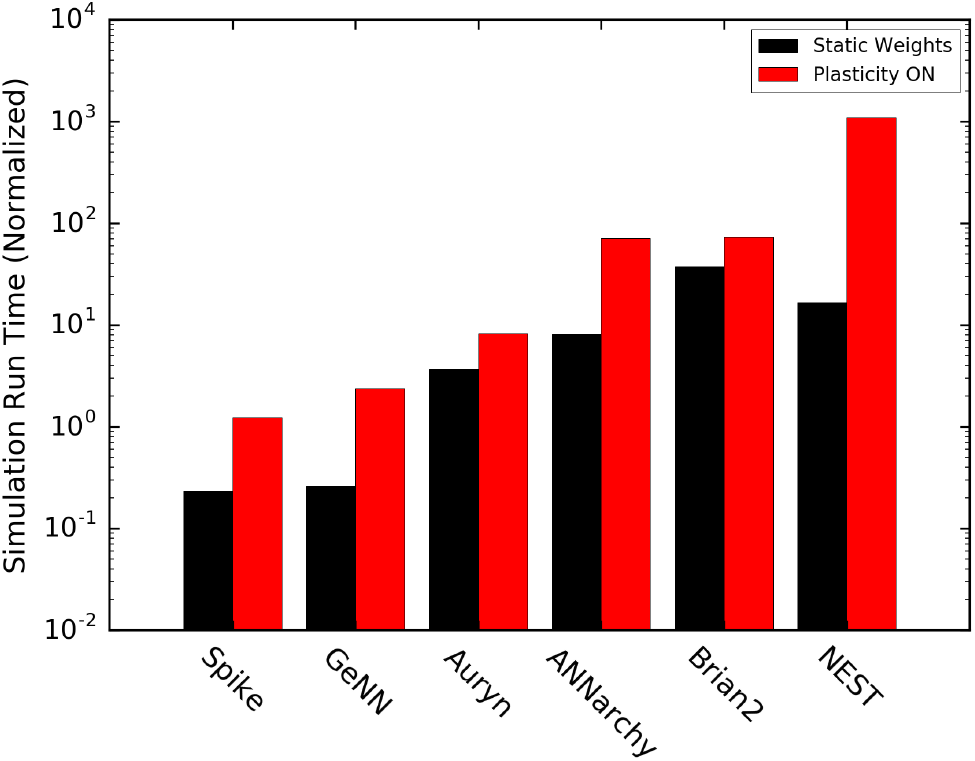
The Brunel benchmark comparison across simulators. All single threaded or single GPU. This benchmark was carried out with and without plasticity as shown in black and red bars respectively. Shorter bars are faster. Note the log scale on the y-axis. These are ordered in speed, fastest first.

The results of simulating in a large network (with and without plasticity) are not qualitatively different to those seen in the prior section (compare Figure 7 to Figure 4). The order of the simulators in terms of mean speed (mean with and without plasticity) is the same. However, the gap between CPU and GPU simulators begins to grow significantly for this larger network. Both Spike and GeNN have a much faster relative simulation time in both the plastic and non-plastic cases and they lead the CPU based simulators in speed. The gap in speed difference between the fastest CPU simulator and the Spike and GeNN simulators has grown in simulations of this scale to an order of magnitude.

In the Vogels-Abbott benchmark, the Spike simulator was approximately four times faster than the Auryn simulator, however in the Brunel benchmark, this rises to approximately seven times faster in the plastic case and approximately 15 times faster in the non plastic case. The speed of the GeNN simulator scales very similarly to Spike and achieves similar performance in these benchmarks (though the plasticity case shows Spike as approximately double the speed of GeNN for this particular setup).

#### 3.2.1 Plasticity with Synaptic Dynamics

The plasticity comparison shown previously (Figure 7) did not show more than a two fold difference in speed between the Spike and GeNN simulators. However, these simulators implement very different approaches to plasticity. The GeNN plasticity benchmark uses an event-based plasticity update mechanism. The code-generation framework employed by GeNN makes this possible through the injection of code relevant to synaptic plasticity into the existing kernels (though a new kernel is also necessary to deal with the arrival of spikes at a post-synaptic neuron). Ultimately, this means that the same code which detects a neuron action potential can immediately alter synaptic weights. By comparison, plasticity in Spike is entirely segregated. All neuron and synapse code for a timestep (or a group of timesteps) is completed before a separate kernel computes any synaptic weight changes that are required. Nonetheless Spike is able to perform competitively.

In order to speed up synaptic plasticity, the GeNN and Spike simulators furthermore avoid computing updates to decaying synaptic traces (see Appendix B.1) on every timestep. Instead, only upon the emission of a spike by the pre or post-synaptic neuron of a plastic synaptic connection are the trace variables updated after taking into account the decay that should have taken place since the previous pre/post-synaptic spike. However, this process of avoiding synaptic updates on every timestep are not possible for more complex synaptic plasticity rules.

To explore the performance benefits of Spike (and in particular TG) in application to plasticity, we consider a more detailed updating of plastic synapses. More complex plasticity rules often rely upon synapse specific dynamics which require a per synapse update upon every timestep. This can effectively make individual synaptic dynamics almost as computationally detailed as the individual neuron membrane voltage dynamics, thereby significantly increasing the computational complexity.

Figure 8 compares the Spike and GeNN simulators with a plastic version of the Brunel benchmark in which synaptic updates (i.e. synaptic traces) are updated every timestep for every synapse. This approximates the computational complexity required for a plasticity rule which requires more detailed synaptic updates. As shown, the speed of the Spike simulator approaches an order of magnitude speed increase over the GeNN simulator when considering detailed synaptic updates. Noteably, this difference in speed is almost entirely accounted for by the timestep grouping (TG) optimisation. Without the inclusion of the TG optimisation, the speed of Spike would be close to that of GeNN. This provides another example of a case in which TG optimisation can provide a significant speedup. It furthermore shows the potential of the Spike simulator in application to plasticity rules which require detailed synaptic dynamics.

**Figure 8.**
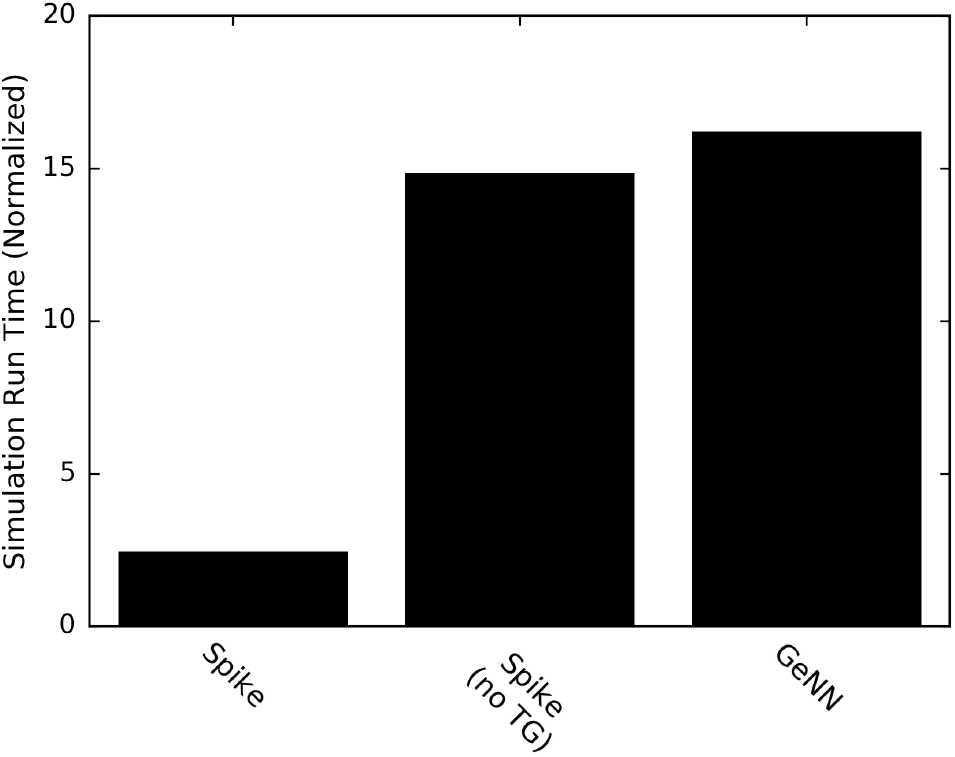
The Brunel benchmark with plasticity via synaptic dynamics. This benchmark was carried out with plasticity active such that every plastic synapse is updated on every timestep. The default Spike simulator is compared to; a case in which timestep grouping (TG) is off, and to the default GeNN simulator. Shorter bars are faster. These are ordered in speed, fastest first.

### 3.3 A Repository of Benchmarks and Comparisons

The production of a set of benchmark comparisons across the set of simulators presented above required an understanding of model construction in these simulators. Despite such comparisons being made previously, code simulating these benchmarks is not widely available. Furthermore, in order to validate that the simulators are producing equivalent network dynamics, a number of tests are also necessary on the outputs of these simulations.

Addressing both of these concerns, we constructed a public repository in which code used to produce the figures throughout this paper is located. This repository includes the compared simulators as submodules and contains a set of iPython notebooks plotting comparisons of network behaviour.

Figure 9 shows two such network behaviour comparisons which are present in the repository. Figure 9 top shows the distribution of Inter-Spike Intervals in the Vogels-Abbott benchmark. This expresses the network firing similarity across simulators. Figure 9 bottom, shows the final weight distribution in the plastic Brunel benchmark with a very similar distribution across simulators. These figures were produced in order to ensure that the networks modelled by the simulators were equivalent. Note that small differences in weight structure in the plastic Brunel benchmark are to be expected given the lack control over the specific input neuron firing times. Specifically, the Brunel network included a set of Poisson firing input neurons and these were implemented separately in each simulator and therefore have different particular spike times.

**Figure 9.**
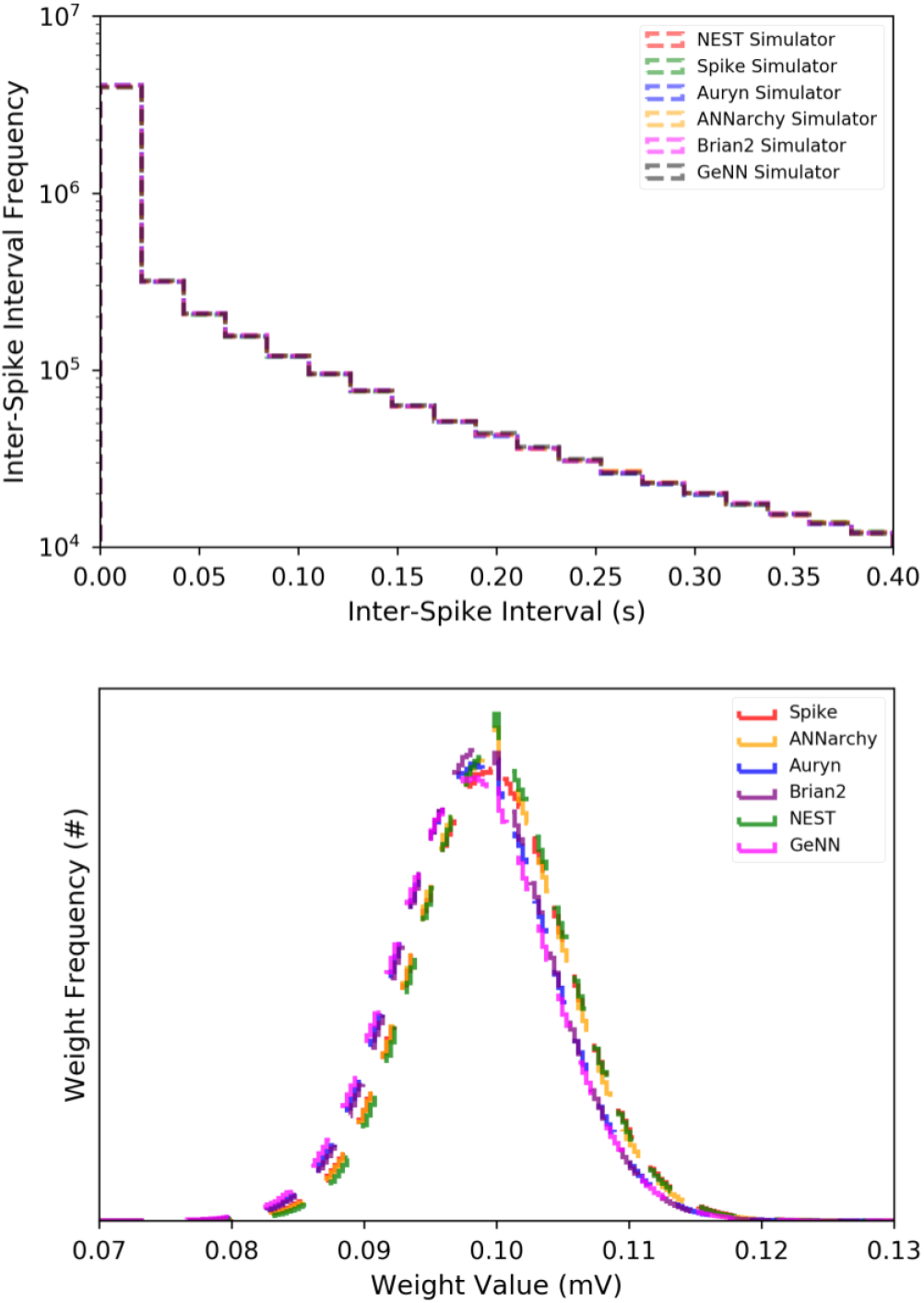
Network properties in the Vogels-Abbott and Brunel benchmarks. Top, the Inter-Spike Interval (ISI) distribution across simulators for the Vogels-Abbott benchmark. Bottom, the weight distribution of the plastic synapses in the Brunel benchmark after 20s simulation time.

## 4 DISCUSSION

The results shown above convincingly place GPU based simulators in the lead for SNN simulation on a single system. As the field moves forward, it is important that such simulators are made easy to use and flexible.

Spike has attempted to make its C++/CUDA form as intuitive as possible to build models in C++. User guides are available and the creation of new models (for neurons, synapses or plasticity rules) is fairly intuitive to someone with a background in C++/CUDA. Nonetheless, writing models in C++ brings with it some complexity given the low-level nature of the code. Furthermore, extensive documentation for Spike has not yet been produced. Given all of this, Spike is placed to act as an simulator with some significant speed benefits and features (in areas such as delay accommodation) over competing simulators, though undoubtedly with some knowledge barriers and inflexiblity in defining new model components.

Though GeNN in its pure C++/CUDA form is a simulator which is fairly difficult to use, the Brian2GeNN project is under development, bringing GeNN as a backend to Brian2. The Brian2 frontend provides a much easier user experience and model definition process. Beyond this, the ability to define models in Python brings major benefits in terms of usability (it being a higher level language).

GeNN’s use of ASG optimisation brings significant benefits to its computational time but it could also potentially integrate optimisations such as TG in order to bridge the speed gap between its performance and Spike’s.

GeNN’s use of code generation furthermore gives it a significant advantage over simulators such as Spike since it is capable of producing tailor-made kernels for a specific simulation. This allows many parameters to be set as constants within kernels and thereby saves time on memory accesses and reduces memory usage. In comparison, Spike’s kernels incur a memory overhead while collecting the specific parameters for individual simulations. On balance, GeNN’s code generation requires a multi-stage compilation process which is not required by Spike. In the most recent version of Brian2GeNN this results in a few seconds of waiting as models are compiled before they are run (though this may well change in the future).

As the field progresses, we expect GPU based simulations to become a common tool for computational neuroscientists. Just as a range of simulators have and continue to exist in the CPU space, we expect the same for GPU based simulators. Code generation approaches, such as those used by GeNN, could prove extremely valuable in producing flexible simulators, whilst more rigid simulations which require high speed and delay insensitivity could turn to Spike. We anticipate further optimisations in the future as GPU based architectures are upgraded and as algorithmic upgrades are proposed.

## 5 CONCLUSION

In order to execute SNN simulations, a range of simulators have emerged with each offering a unique combination of speed, hardware integration, and ease of model definition. In this study we compared the ANNarchy (Vitay et al., 2015), Auryn (Zenke and Gerstner, 2014), Brian2 (Stimberg et al., 2014), GeNN (Yavuz et al., 2016), and NEST (Linssen et al., 2018) simulators against the Spike simulator. These comparisons showed the efficacies of GPU based SNN simulators of which GeNN and Spike are examples.

We successfully showed a significant speed benefit with the use of GPU based simulators. These comparisons were carried out both on single-threaded systems and in a multithreaded case where CPU based simulators were shown to have insufficient speed to outperform GPU based simulators. In the case of larger networks, the difference in GPU simulation time to CPU simulation time exceeded an order of magnitude and we therefore identify GPUs as a step forward for point-neuron based SNN simulations.

Moving forward, more flexible simulators such as GeNN are expected to be invaluable in allowing both ease of model definition and speed of simulation. Spike is an alternative simulator which we show has a significant speedup over other simulators and is “delay insensitive” – allowing heterogeneous synaptic delays without any change in simulation speed. In order to ensure continued comparison and benchmarking of SNN similators, we publish a repository which contains all of the code used to produce the graphs in this paper and more.

## AUTHOR CONTRIBUTIONS

NA was the original creator of Spike. NA conceived of and developed the contents of this paper (including all figures). JBI and TSS are core members of the development team in Spike and have heavily affected the form that the Spike software currently takes. SMS supervises NA, JBI, and TSS.

NA wrote this paper and it has been contributed to by SMS, JBI and TSS.

## ACKNOWLEDGMENTS

Friedemann Zenke (creator of the Auryn simulator (Zenke and Gerstner, 2014)) has been an unmatched source of advice and information through the production and development of the Spike simulator. Personal thanks from NA for the many discussions and emails.

## DATA AVAILABILITY STATEMENT

The Spike Simulator is available at;

https://github.com/OFTNAI/Spike

The benchmarks, simulation code and timing data for this study can be found in the repository;

[SNNSimulatorComparison]
[https://tinyurl.com/y7gltmsw]

## APPENDICES

### A Vogels-Abbott Benchmark Model

The Vogels-Abbott benchmark is based upon a reduced scale version of the network presented in the Vogels and Abbott (2005)
publication. This reduced scale version is detailed in the paper by Zenke and Gerstner (2014).

The network consists of 3200 excitatory and 800 inhibitory Leaky Integrate and Fire (LIF) neurons. Synaptic connections influence a conductance based current input model and the network is driven by constant background stimulation. Individual neuron dynamics can be described;

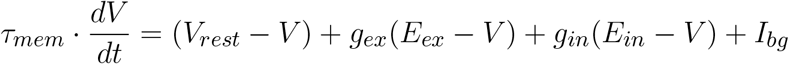

where *V* is the neuron membrane voltage, *τ_mem_* is the membrane time constant (which can also be computed by dividing the membrane capacitance by the cell leakage conductance) and *V_rest_* is the cell resting potential. Synaptic inputs are governed by the excitatory and inhibitory conductances, *g_ex_* and *g_in_* respectively, and the reversal potentials, *E_ex_* and *E_in_* respectively. Finally, cells are stimulated by a constant input *I_bg_*.

If the membrane potential of a cell reaches a threshold *V_thresh_*, it emits an action potential (or spike) and is thereafter brought back to the reset potential *V_reset_* for a period of time equal to the refractory period of the cell *τ_ref_*. After this refractory period, the cell dynamics are allowed to continue.

Synaptic connections update the excitatory and inhibitory synaptic conductances. A pre-synaptic spike on a synaptic connection causes a discontinous jump in the corresponding post-synaptic cell synaptic conductance. For excitatory synaptic connections, a pre-synaptic spike causes a jump in the post-synaptic neuron excitatory synaptic conductance, after a time dictated by the axonal delay, such that *g_ex_* ← *g_ex_* +*w_ex_* where *w_ex_* is the weight of the excitatory synaptic connection. Similarly, inhibitory synaptic connections cause a discontinous jump in the inhibitory synaptic conductances of post-synaptic cells such that *g_in_* ← *g_in_* + *w_in_* where *w_in_* is the weight of the synaptic connection. Finally, the cell synaptic conductances undergo dynamics;

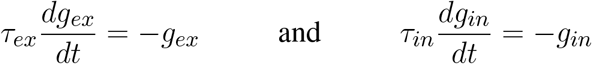

The parameters used for this model are detailed in Table 1.

**Table 1.**
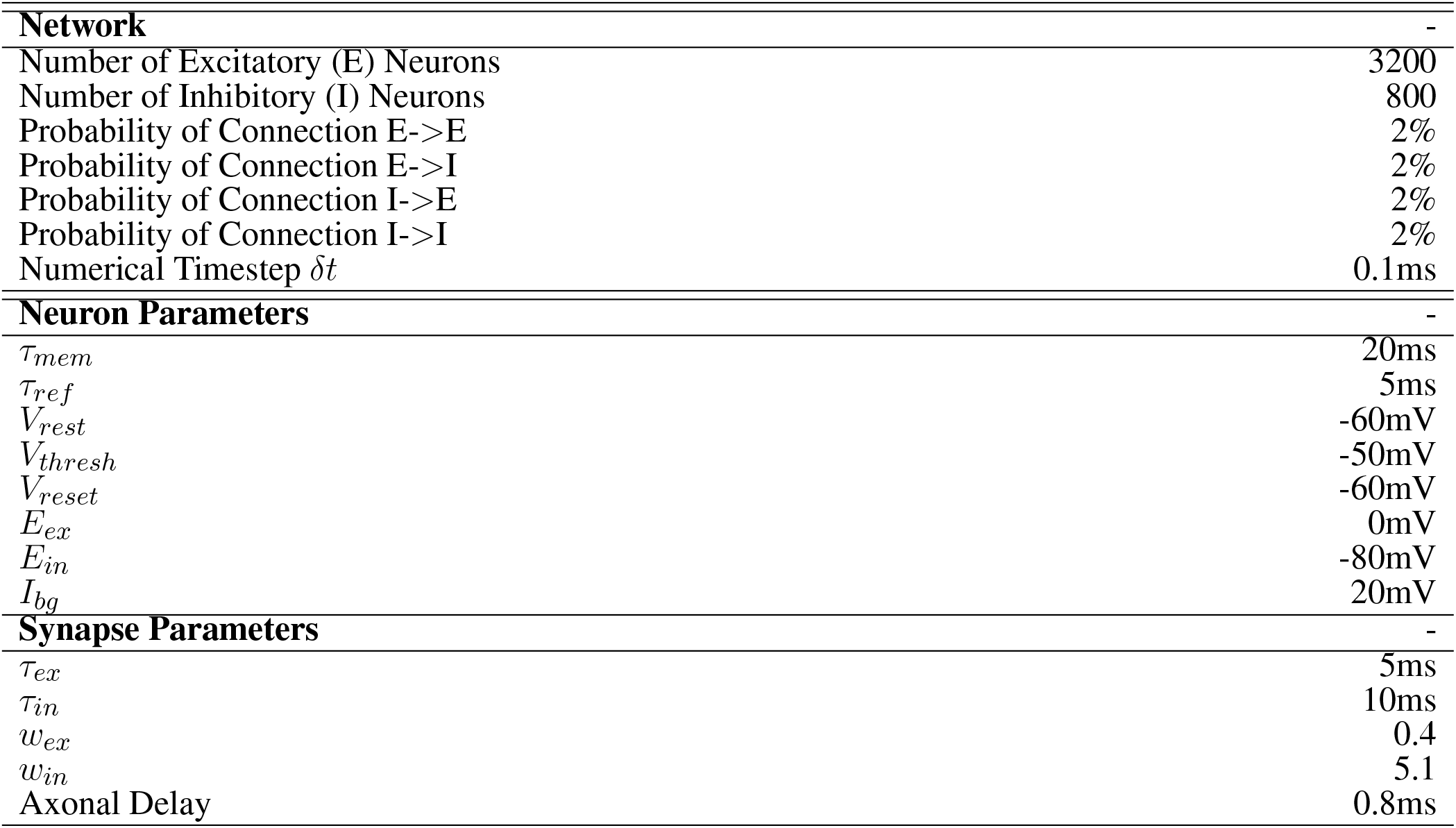
Network, Neuron, and Synaptic Parameters used for the Vogels-Abbott Benchmark.

|  |  |
| --- | --- |
| **Network** | - |
| Number of Excitatory (E) Neurons | 3200 |
| Number of Inhibitory (I) Neurons | 800 |
| Probability of Connection E->E | 2% |
| Probability of Connection E->I | 2% |
| Probability of Connection I->E | 2% |
| Probability of Connection I->I | 2% |
| Numerical Timestep $\delta t$ | 0.1ms |
| **Neuron Parameters** | - |
| $\tau_{mem}$ | 20ms |
| $\tau_{ref}$ | 5ms |
| $V_{rest}$ | -60mV |
| $V_{thresh}$ | -50mV |
| $V_{reset}$ | -60mV |
| $E_{ex}$ | 0mV |
| $E_{in}$ | -80mV |
| $I_{bg}$ | 20mV |
| **Synapse Parameters** | - |
| $\tau_{ex}$ | 5ms |
| $\tau_{in}$ | 10ms |
| $w_{ex}$ | 0.4 |
| $w_{in}$ | 5.1 |
| Axonal Delay | 0.8ms |

### B Brunel Benchmark Model

The Brunel benchmark is based upon a network adapted by Zenke and Gerstner (2014), published in its original form by Brunel (2000).

The network consists of 8000 excitatory and 2000 inhibitory Leaky Integrate and Fire (LIF) neurons. Synaptic connections influence a voltage injection based input model and the network is driven by 10,000 excitatory input neurons with 20Hz random Poisson firing. Individual neuron dynamics can be described;

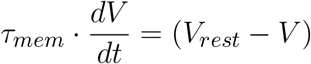

where *V* is the neuron membrane voltage, *τ_mem_* is the membrane time constant (which can also be computed by dividing the membrane capacitance by the cell leakage conductance) and *V_rest_* is the cell resting potential. Synaptic inputs are governed by the excitatory and inhibitory voltage injections. For excitatory or inhibitory synaptic connections, following a pre-synaptic neuron spike (and after awaiting any axonal delay), the voltage is directly modified in a discontinuous fashion such that *V* ← *V* + *w* where *w* is the weight of the synaptic connection. All excitatory synaptic connections are initialised with a weight *w_ex_* and all inhibitory synaptic connections are initialised with a weight *w_in_*.

If the membrane potential of a cell reaches a threshold *V_thresh_*, it emits an action potential (or spike) and is thereafter brought back to the reset potential *V_reset_* for a period of time equal to the refractory period of the cell *τ_ref_*. After this refractory period, the cell dynamics are allowed to continue.

The parameters used for this model are detailed in the Table 2.

**Table 2.** Network, Neuron, and Synaptic Parameters used for the Brunel Benchmarks.

|  |  |
| --- | --- |
| **Network** | - |
| Number of Excitatory (E) Neurons | 8000 |
| Number of Inhibitory (I) Neurons | 2000 |
| Number of Excitatory Poisson Input Neurons (P) | 10000 |
| Probability of Connection E->E | 10% |
| Probability of Connection E->I | 10% |
| Probability of Connection I->E | 10% |
| Probability of Connection I->I | 10% |
| Probability of Connection P->E | 10% |
| Probability of Connection P->I | 10% |
| Numerical Timestep $\delta t$ | 0.1ms |
| **Neuron Parameters** | - |
| $\tau_{mem}$ | 20ms |
| $\tau_{ref}$ | 2ms |
| $V_{rest}$ | 0mV |
| $V_{thresh}$ | 20mV |
| $V_{reset}$ | 0mV |
| Poisson Neuron Firing Rate | 20Hz |
| **Synapse Parameters** | - |
| $w_{ex}$ | 0.1mV |
| $w_{in}$ | -0.5mV |
| Axonal Delay | 1.5ms |

#### B.1 Plasticity

The Brunel benchmark can be run with or without a plasticity rule active. When active, the plasticity rule is a Spike-Timing Dependent Plasticity (STDP) rule which updates synaptic connections upon pre and post-synaptic neuron action potentials.

The STDP rule applied in this model is a weight dependent STDP rule such that for each synapse, we have two traces – one for its pre-synaptic neuron and one for its post-synaptic neuron, *z_pre_* and *z_post_*. Upon a pre-synaptic spike, we wait for a duration equal to the axonal delay, following which we update the pre-synaptic trace such that; *z_pre_* ← *z_pre_* + 1.0. Upon post-synaptic spikes, the post-synaptic trace is updated such that; *z_post_* ← *z_post_* + 1.0. These traces both decay with the dynamics that follow.

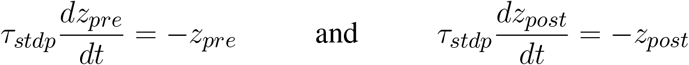

Long Term Depression (LTD) is implemented as follows. Upon a pre-synaptic spike (after awaiting the axonal delay), the weight *w* of the synaptic connection is also updated;

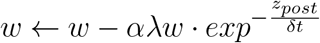

where *λ* is the learning rate, and *α* is a scaling factor which is only applied to LTD (thus dictating the relative strength of LTD).

Long Term Potentiation (LTP) is implemented as follows. Upon a post-synaptic spike, the weight *w* of the synaptic connection is also updated;

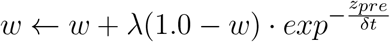

The plastic synaptic connections also have a hard bound upon their weight such that if the weight *w* increases above a maximum weight value *w_max_*, it is bounded and *w* ← *w_max_*. Similarly, if the weight *w* reduces below zero, it is bound and set to zero; *w ←* 0.0.

This STDP rule is only applied to the excitatory to excitatory (*E*–> *E*) synaptic connections in this model. All parameter values are presented in Table 3.

**Table 3.** Network, Neuron, and Synaptic Parameters used for the Brunel Benchmarks.

| Plasticity Parameters |  | - |
| --- | --- | --- |
| $\tau_{stdp}$ | | 20ms |
| $\alpha$ | | 2.02 |
| $\lambda$ | | 0.01 |
| $w_{max}$ | | 0.3mV |

